# Same place, different time: Leopards employ temporal partitioning as a co-existence mechanism in a human-dominated, unprotected agricultural landscape in Sri Lanka

**DOI:** 10.1101/2025.10.10.681667

**Authors:** Andrew M. Kittle, Anjali C. Watson

**Affiliations:** The Wilderness & Wildlife Conservation Trust, Colombo, Sri Lanka

**Author notes:** corresponding author: (AMK).

## Abstract

As humans and wildlife increasingly overlap on a rapidly changing planet, it is important to understand co-existence mechanisms in unprotected landscapes, especially for large carnivores with which humans share an uneasy history. In Sri Lanka’s tea estate-dominated Central Highlands, a dense, widespread human population shares the landscape with the island’s apex predator, the Sri Lankan leopard. This study used remote cameras to determine leopard density, generalized linear models to investigate leopard space-use in relation to key anthropogenic, ecological and environmental variables, and fitted kernel density curves to identify temporal activity patterns in this unprotected, human-dominated region. Adult leopard density (8.82/100 km²) was lower here than in key Sri Lankan National Parks, but not significantly so, suggesting good leopard habitat suitability. Spatially, leopards did not avoid humans but incorporated anthropogenic factors into space-use decisions, with areas of intermediate human and dog presence disproportionately used. Male leopards were more likely to occur at intermediate distances from human settlements, perhaps balancing risk with efficiency while navigating this human-dominated landscape. Female leopards biased space-use towards areas of high natural prey availability, aligning with their need to provide for cubs. Temporally, leopards were significantly less diurnal and more active at dusk than their protected counterparts, likely due to temporal partitioning from diurnally active humans, an adaptation which allows for human-leopard co-existence. This is the first comprehensive study of leopards outside protected areas in Sri Lanka, and highlights the value of unprotected landscapes and their importance for long-term leopard conservation. To foster human-leopard co-existence it is necessary to maintain current human activity patterns which allow for temporal partitioning; ensure existing natural prey resources remain to prevent leopards switching to domestic prey species; and reduce/eliminate the use of wire snares which transcend the effect of temporal partitioning by de-coupling persecution risk from human presence.

## Introduction

Protected areas (PAs) represent vital refuges for wildlife and are key components of long term global conservation efforts [1,2,3]. However, the vast majority (∼85%) of the world’s terrestrial area is unprotected [4], with wildlife great and small – from amphibians [5] to birds [6] to large mammals [7] relying – wholly or in part - on these unprotected landscapes for their survival, either for migration [8], dispersal [9], foraging [10], or simply as part of their daily range [11]. It is widely accepted that unprotected landscapes pose greater risks to wildlife than PAs, mostly due to their widespread use by humans and the land use changes they affect [12]. Relatively little is understood, however, about how wildlife utilize these typically human-dominated landscapes, especially in relation to what is known about their behavior and ecology within PAs. With the world’s human population and ecological footprint continuing to grow [13], and climate change impacts expected to increasingly see wildlife species move out of their traditional regions [14], it is imperative to better understand how they use shared spaces.

For larger species - especially top carnivores which have a long and complex relationship with humans [15], are often viewed negatively by humans [16,17] and are therefore targets of persecution [18] - understanding the mechanisms by which co-existence with humans can be promulgated is key to their long-term viability. Top carnivores can play vital roles within ecosystems and their disappearance can have cascading effects across trophic levels, yet these large predators are being extirpated at an alarming rate [18]. Just as the study of human-wildlife conflict has shed light onto important factors that create tension between people and the animals with which they share space [19], it is increasingly accepted that equal consideration must be given to understanding the factors that promote human-wildlife co-existence so that these factors, that allow people and wild animals to successfully share space, can be incorporated into wildlife policy, planning and management [20].

Unprotected areas are often diminished in terms of natural prey availability [11], which is a key determinant of large predator home range size and therefore density [21]. Furthermore, these areas tend to be associated with increased risk [22] and often have sizeable areas – e.g. city centres, agricultural expanses - that are not compatible with large carnivore use, meaning individuals require larger areas in which to accomplish basic activities such as foraging, mating and rearing young. As such, it might be expected that carnivore densities outside PAs are lower than those found within comparable, well-protected PAs. A key initial consideration for the management of wildlife in unprotected areas is to understand how many reside there.

Spatial partitioning between humans and wildlife has long been seen as a key concept of successful, broad-scale co-existence; it is this concept that underlies the establishment and maintenance of Protected Areas – places for wildlife distinct from places for people [23]. However wild animals do not respect artificial boundaries and human populations continue to expand into new areas, so in much of the world humans and wildlife do share space [24]. Even outside PAs, apex predators often employ fine scale spatial avoidance of humans and/or human infrastructure, such as the avoidance by lynx (*Lynx lynx*) of dense road networks within their home ranges in Norway [25], the avoidance of human settlements by tigers (*Panthera tigris*) in China [26] or simply the avoidance of areas of high human presence by lions (*Panthera leo*) [27]. In some landscapes, however, spatial partitioning is particularly challenging due to high human densities or a widespread human footprint across space [13], and here it is increasingly seen that top predators employ temporal partitioning to avoid encountering people [28,29].

Sri Lanka is a relatively small (65,610 km^2^), densely populated (336/km^2^), mostly rural (∼80%) island that is – together with India’s Western Ghats - one of the world’s 36 global biodiversity hotspots [30]. In Sri Lanka, the leopard (*Panthera pardus kotiya*) is a threatened, endemic sub-species and the island’s apex predator [31]. Despite a low estimated population of < 1000 mature individuals [32], leopards in Sri Lanka remain extant in all climatic zones and are present in both protected and unprotected areas [31].

In Sri Lanka’s Central Highlands – a UNESCO World Heritage site based on its high levels of biodiversity and species endemism [33] – leopards reside in and around the unprotected, tea estate-dominated landscape [34] where every year several animals are killed due to human persecution [35]. Prior to the widespread clearance of forest for the cultivation of first coffee (1830 - 1880) and then tea (*Camillia sinensis*; 1880 - present) during British colonial times, the Central Highlands were blanketed in thick sub-montane and montane forest [36], of which mostly small, isolated patches remain. Despite anecdotal tales of leopards in and around these forest patches from colonial days to the present, and despite research on the leopard being conducted in the region’s most iconic PA, Horton Plains NP [37,38], almost nothing is known about the behavior and ecology of leopards in these unprotected tea estate landscapes (but see [34]).

The objective of this study was therefore to improve our understanding both of leopard ecology and behavior outside protected areas in Sri Lanka and of how these apex predators have adapted to sharing space with humans. Specifically, we wanted to test three hypotheses related to human-wildlife co-existence: 1) Human-dominated landscapes are sub-optimal for leopards. If this is supported, we would expect to see leopard’s living at a lower density in this landscape compared to within the island’s foremost protected areas. 2) Leopards employ spatial partitioning to reduce threats from humans. If this is supported, we would expect to see leopards avoiding areas of high human use and/or infrastructure. 3) Leopards employ temporal partitioning to reduce threats from humans. If this is supported, we would expect to see less diurnal activity in this unprotected, heavily cultivated landscape compared to protected ones on the island.

## Materials and Methods

### Study area

This study was conducted in the southern region of the Central Highlands of Sri Lanka, in the Upper Kelani Valley catchment area surrounding the Moussekelle reservoir, immediately north of the Peak Wilderness Sanctuary (Fig. 1). The region is characterized by vast expanses of tea cultivation, interspersed with fragmented, often secondary forest patches, dense shrub patches where tea has been “released”, and Eucalyptus (*Eucalyptus sp.)* and Pine (*Pinus caribaea*) plantations [39]. The region is within Sri Lanka’s sub-montane wet zone with annual rainfall > 2500 mm. Although it rains on more than half the days across all months, rainfall is heaviest in October and November (> 300 mm/month) when the area is impacted by the Northeast monsoon [40]. Rainfall is also heavy in April and May (250 – 300 mm/month), roughly coinciding with the southwest monsoon (May – Sept). The dry season (< 150 mm/month) runs from January through March. Average temperatures range from 18° - 30°C, although on the upper slopes nighttime temperatures can dip to 12°C [40]. Elevation in the study area ranges from 1000 – 1800 MASL, with steep ridgelines running between shallow, flattened valleys where rivers and streams flow.

**Fig. 1:**
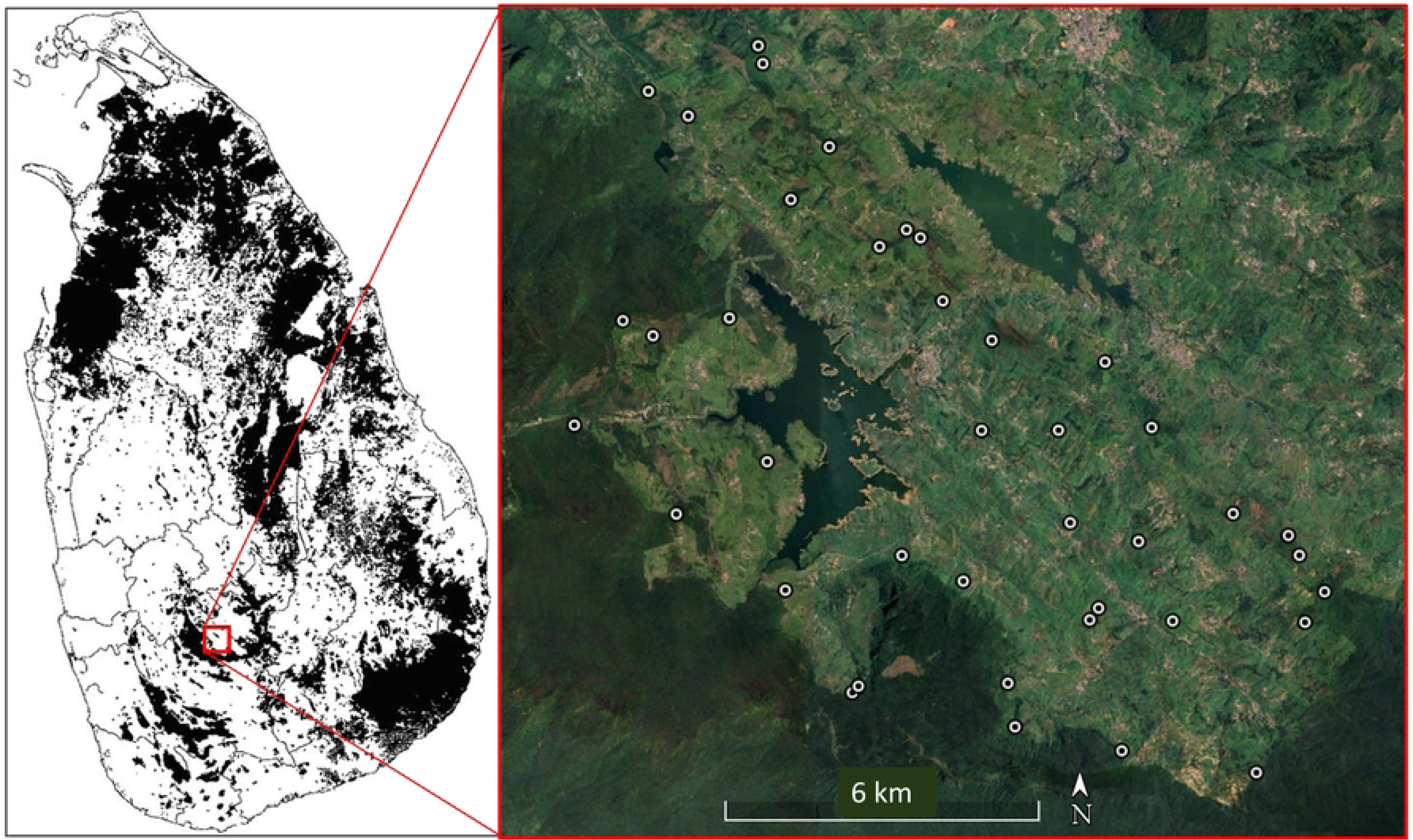
Map of Sri Lanka showing current forest cover (black shading) with the inset showing the study area. The white dots are remote camera locations (N = 40) utilized during this study. The two large hydro-electric reservoirs can be seen with remote camera locations surrounding the Moussekelle reservoir and to the south west of the Castlereagh reservoir. Dark green along the southern and western regions is the Peak Wilderness Protected Area complex with the lighter green being the typical mosaic of tea cultivation, small remnant forest patches, plantations, grasslands and communities.

In this region of the Central Highlands the tea industry is dominated by large Regional Plantation Companies, each of which typically runs several neighbouring estates. Each estate is divided into several (∼2-8) Divisions with each Division typically holding at least one community comprised of estate workers and their families. Larger towns are also widespread, typically along valley bottoms where waterways and main roads are also located. As such, the human footprint in this landscape is large and extensive. The human population density of the Nuwara Eliya District in which the study area is located is > 400/km² [41].

### Leopard density

To estimate leopard density, we set 40 remote camera stations throughout the Central Highlands landscape surrounding the Moussekelle reservoir from August - October 2016 (Fig. 1; Table S1). Since adult leopards across continents and landscapes frequent roads and other linear pathways while patrolling their territories [37,42,43,44,45], cameras were set adjacent to and facing unpaved tea estate roads, walking paths or animal trails. Twenty of the stations comprised two cameras, one either side of these trails, with the balance 20 stations having a single remote camera. There was some variation in the distance from trails (1 – 3m) of cameras and camera height (30 cm – 1 m) due to the heavily cultivated landscape which necessitated use of infrequent trees or sturdy tea bushes, but cameras were nevertheless set to ensure photo-capture of leopards and their main prey [46].

The study was divided into 3 rounds, with round 1 employing 12 camera locations across 36 days, round 2 using 14 camera locations across 35 days and round 3, 14 camera locations across 44 days (Table S1). The un-buffered camera array covered ∼ 140 km² (Fig. 1) with locations spaced an average of 1.2 km apart, to ensure all prospective leopards were exposed to detection in this unknown landscape and optimize the trade-off between area coverage and photographic recaptures while ensuring a closed population [46]. To ensure sufficient image quality for accurate identification of individuals, we used Scoutguard SG565F (Boly Inc., Shenzen, China) incandescent flash cameras with sensitivity set to “normal” and minimal camera delay (“0”) to ensure capture of multiple animals moving together.

Photo-captured leopards were individually identified using their unique spot patterns, with age and sex categorized based on morphological differences [45]. For each individual a capture history was developed but only mature leopards (estimated to be ≥ 2 years old) were used for population density analysis. Both flanks of mature leopards were almost all clearly photo-captured, except one adult female whose left flank only was photographed and three unsexed individuals whose right flanks only were photographed. Since the three unsexed individuals were not clear images, we elected to utilize only individuals who had been photographed on the left flank.

We used spatially explicit capture-recapture (SECR) models within a maximum likelihood (ML) framework (*secr* 4.6.0 in R) [47] to estimate leopard density. The SECR process, which fits spatial models of the population and detection process to individual spatial detection histories, is considered unbiased by edge effects or incomplete detection [48]. SECR models were constructed using count detectors to account for multiple or repeat individuals captured by each trap on each occasion. Given the variation in the number of days that individual camera stations were active, we incorporated varying effort by valuing complete 24-hour camera occasions as 2, 12-hour occasions as 1, and inactive days as 0 [49]. We fitted a half-normal detection function with Poisson distribution, which is the most commonly employed function in spatial capture-recapture analysis [50]. This function describes the probability of capture (*P*) of individual *i* at trap *j* as a function of the distance (*d*) from that individual’s activity centre to the trap using the equation *Pij* = *g0* exp (-*dij*^2^/2σ ^2^)), with *g0* the probability of capture at home range centre and *σ* a spatial parameter related to home range size [50]. Since our study area is centred around two large hydro-electric reservoirs (Fig. 1), we created a habitat mask with a 15km buffer around the remote camera array and with the reservoirs defined as “non-habitat” and removed. We did not consider any other areas “non-habitat” as leopards in this heavily fragmented landscapes are known to occasionally be seen even in human communities. We used *secr*’s “suggest.buffer” function to determine a suitable starting width (7601m) for the concave polygon buffer around the detector array and then, using the “make.mask” function, created a set of habitat masks starting with a smaller buffer (4000m) and sequentially increasing its width, by 1000m, around the trapping grid, up to 15000m. To ensure that individuals outside the buffer area cannot be captured within and artificially inflate density estimates, we then used the “check.mask” function to compare models using the 12 mask size iterations until model LogLikelihood estimates stabilized at 3 decimal places (Table S2) [48,51] and utilized that buffer size for further analysis.

Variation between sexes in both range size and use structures leopard spatial dynamics [52] and male leopards in Sri Lanka typically utilize much larger areas than females [45,53,54]. Therefore, we fit 2-class hybrid mixture models (hcov in *secr*) with sex modelled as a covariate [51,53,55] allowing for varying capture parameters (*g0* and *σ*) between sexes. Using Akaike’s Information Criterion for small sample sizes (AIC*c*) as well as Akaike weights (AIC*cwt*) [56] to compare them, we tested 4 models: 1) a null model without variation in capture parameters; 2) a model allowing sex-specific variation in *g0*; 3) a model allowing sex-specific variation in *σ*, and 4) a model allowing sex-specific variation in both capture parameters.

### Leopard site utilization

At each of the 40 camera stations, as the dependent variable, we determined the total number of leopard observations, inclusive of all age/sex classes, which were considered to be independent if >1 hour passed between detections of the same individual at the same site. If leopard detections <1 hour apart were of different individuals– based on age, sex and/or individual identification – these were also utilized for analysis.

To investigate the factors driving occurrence patterns, each location was characterized by ecological and anthropogenic attributes selected to represent one of four specific underlying factors: human-leopard co-existence, habitat suitability, landscape topography and prey availability. The human-leopard co-existence variables were 1) the relative abundance index of humans at each camera location (RAI = (# of human or vehicle detections/# remote camera 24-hour surveillance periods) * 100), 2) the RAI of domestic dogs, 3) the distance (m) to nearest human settlement, and 4) the distance (m) to nearest road. Distance to human settlement was determined using the “distance measure” tool on Google Earth as available GIS layers did not have sufficiently fine grain to identify all communities on the landscape. Distance to road employed GIS shape files acquired from the Sri Lanka Survey Department [57] of paved primary (Main road A Grade) and secondary (Minor road B Grade) roads with the distance determined using the “Near” tool under “Proximity” analysis in ArcGIS. Habitat suitability metrics were based on the importance of protected areas (PAs) and forest cover for leopard distribution in Sri Lanka [31]. Distance to PA was determined in the same way as distance to roads above, only utilizing the Sri Lanka Protected Area layer [57]. Distance to forest was determined in the same way as distance to settlement above, since again, the grain of available GIS layers was not sufficiently fine to allow identification of small forest patches in this landscape. For this analysis, forests were considered all treed landscapes inclusive of plantation forests as leopards are generalists [58] and these habitats are used by leopards, especially in landscape mosaics [59]. Landscape topography was represented by a measure of elevation (MASL) taken with a hand-held GPS unit at each remote camera location. Prey availability was represented by black-naped hare (*Lepus nigricollis*) RAI and red muntjak (*Muntiacus muntjak*) RAI as both species are widespread in the tea estate landscape and important prey species here [60], with the former the most available on the landscape [60] and the latter the only potential prey species whose weight is within the range “highly preferred” by leopards globally (23 – 25 kg) [61]. The original variable list contained two metrics of available prey biomass – total biomass and biomass of species within the weight range (10 – 40 kg) of leopard preference [61] – however correlation analysis revealed these variables to be highly correlated (r > |0.8|) with each other and several other independent variables, so were dropped from analysis. The final list of independent variables all had correlation coefficients < |0.7| (Table S3). Since some relationships might be non-linear, with leopards more likely to be found at moderate levels of a given variable [62], we employed the quadratic of four independent variables: Human_RAI, Dog_RAI, Distance to Settlement and Elevation (Table S3).

The number of remote camera stations (40) dictated we keep the analysis simple according to Harrell’s rule, whereby each variable should have ∼10 observations for meaningful interpretation [63,64]. As such, the most complex model contained only 4 independent variables. Due to the preponderance of 0s in the dataset, we utilized Generalized Linear Models (GLMs) with a Poisson distribution. Since this method requires the dependent variable to be integers, we could not use rates, so instead employed the total number of independent detections at each remote camera site (as detailed above) and incorporated an offset of the log of the number of 24-hr periods the cameras were active, into the models to ensure that survey effort was accounted for. For ease of interpretation and comparison we scaled all independent variables using the “scale()” function in R so that each had a mean of 0 and standard deviation of 1. The same set of models was used for all leopards combined as well as for male and female leopards separately, in order to understand sex-based variation in habitat suitability.

We determined Akaike’s Information Criterion for small sample sizes (AIC*c*) for each model and used ΔAIC*c* and model weights (*w_i_*) to rank models [56]. Model adequacy was determined using standard analyses of residuals (*i.e*. residuals vs fitted values plots, normal Q-Q plots, scale-location plots, and residuals vs leverage plots). Model goodness-of-fit was determined using McFadden’s psuedo-R² (1 – (residual deviance / null deviance)).

### Leopard activity times

All photo-captures recorded the date and time of capture, and these times – after conversion to radians - were used to create fitted kernel density curves using default smoothing parameters in R package “overlap” [65]. A primary goal was to compare the activity patterns of leopards in the unprotected tea landscape to those in established protected areas in Sri Lanka, so we utilized existing remote camera data from Wilpattu National Park (WNP) in July to October 2015 [45] and Gal Oya National Park (GONP) from November 2017 – June 2020 (Table S4). WNP is Sri Lanka’s largest PA (1317 km²), located in northwestern Sri Lanka’s arid and dry zones, whereas GONP (259 km²) is in the Intermediate zone and protects the watershed forest of Sri Lanka’s largest reservoir, the Senanayake Samudra. In both locations white flash remote cameras (Scoutguard in WNP; Cuddeback and Scoutguard in GONP) were set up along jeep tracks, walking paths and animal trails in a similar manner to the unprotected tea estate landscape. Camera height placement and settings were the same across sites. Leopard photo-captures for these sites were also used to create fitted kernel density curves as above.

In order to categorize leopard activity, the 24-hrs that characterize a full day were divided into discrete periods based on sunrise and sunset times in Sri Lanka [66] (Table S5). Across the year sunrise ranged from 5:48 – 6:26 and sunset from 17:45 – 18:28. To determine crepuscular periods we therefore created a buffer of ∼30 minutes of these minimum and maximum times, with diurnal and nocturnal periods therefore the remaining hours of light and dark, respectively, between crepuscular periods. Final time periods were defined as crepuscular (dawn) from 5:15 – 7:00, diurnal from 7:00 – 17:15, crepuscular (dusk) from 17:15 – 19:00, and nocturnal from 19:00 – 5:15.

To categorize leopard activity in the three locations we calculated Manly selectivity measures using ratios of use vs. availability for each time period [67]. The equation used was:

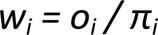

where *w_i_* is the selection ratio for the period *i*; *o_i_* is the proportion of leopard trap events (photo-captures) in period *i*; and *π_i_* is the proportion of length of time in period *i* to the length of time in all periods. When *w_i_* > 1 the time period *i* is considered selectively used (*i.e.* use > availability) and conversely, when *w_i_* < 1 the time period *i* is considered avoided (*i.e.* availability > use; [67,68,69].

To compare leopard activity patterns between the protected (WNP and GONP) and unprotected (tea estate landscape) areas, we compared the independent proportions of leopard observations for each of the time periods between all sites. To do this we employed a two-sample, two-tailed Z-test [70].

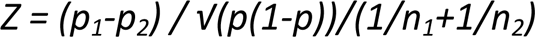

where p_1_ and p_2_ are the sample proportions, n_1_ and n_2_ are the sample sizes, and p is the total pooled proportion calculated as:

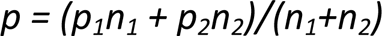

We used p-values at a significance level of 0.05 to determine effect and whether the z-value was positive or negative to determine the direction of effect. We also calculated 95% confidence intervals (CI) for each effect with the interpretation that CIs that do not overlap 0 represent strong effects.

## Results

Across 1258 24-hour remote camera periods we recorded 90 separate leopard detections, 94.4% of which were definitively individually identified (N=85). Three leopard images were not clear enough to determine age/sex or individual ID, while 2 additional photo captures of adult males could not be individually identified with certainty. These images were discarded from the density analysis, as were the two cub images, resulting in 83 leopard photo-captures used for density analysis. The leopard site use analysis used all 90 leopard images, with 54 photo-captures of adult males used for the “male only” site use analysis and 31 photo-captures of adult females used for the “female only” site use analysis. In total, 18 individual leopards were identified, 16 mature individuals (6 M, 10 F) and 2 cubs. Leopards were photo-captured at 26 (65%) camera stations.

### Leopard density

The top-ranked model (AIC*cwt* = 0.904) for estimating leopard density allowed for heterogeneity in the spatial parameter related to home range size (*σ ∼ h2*; Table 1). Density estimates stabilized with a buffer width of 11000m (Table S2), so employing the above hybrid-mix model formulation with varying effort and 11000m buffer, we estimated the study area’s mature leopard population density as 8.82 ± SE 2.42 leopards/100 km² (95% CI = 5.21 – 14.95). Adult sex ratio (ASR) of observed individuals (37.5%; 1M: 1.67F) was considerably different from that determined by the hybrid mixture model (*pmix*) (16.1%; 1M: 5.17F; Table 2). The probability of detection at home range centre (*g0*) was 0.046 ± SE 0.008 for both sexes, whereas the spatial parameter (*σ*) was 2409m ± SE 240 for males and 737m ± SE 87 for females (Table 2).

**Table 1:**
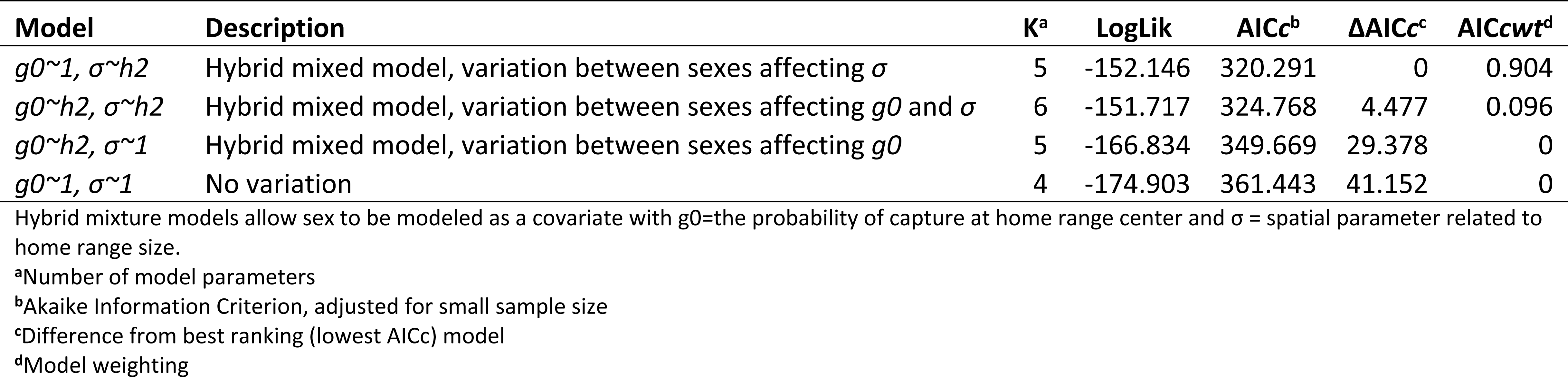
Comparison of spatially explicit capture-recapture (R package *secr*) model selection parameters for 2-class hybrid mixture models (*hcov* in R).

**Table 2:** Parameters estimated by the best spatially explicit capture-recapture (R package *secr*) model for leopard density estimation (g0∼1, *σ∼h2*).

| Parameter<sup>a</sup> | Mean±SE | | 95% CI | |
| --- | --- | --- | --- | --- |
|  | ♂ | ♀ | ♂ | ♀ |
| *g0* | $0.046 \pm 0.008$ | $0.046 \pm 0.008$ | 0.032 - 0.065 | 0.032 - 0.065 |
| $\sigma$ | $2409 \pm 240.1$ | $737 \pm 87.0$ | 1983 - 2927 | 585 - 928 |
| *pmix* | $0.161 \pm 0.073$ | $0.839 \pm 0.073$ | 0.063 - 0.357 | 0.643 - 0.937 |

<sup>a</sup>*g0* = probability of capture at home range centre; $\sigma$ = spatial parameter (in meters) related to home range size; *pmix* = sex ratio

The observed density was lower than what has been found in Sri Lanka’s best protected National Parks, but in most cases this difference is not significant as evidenced by overlapping 95% CIs (Table 3).

**Table 3:**
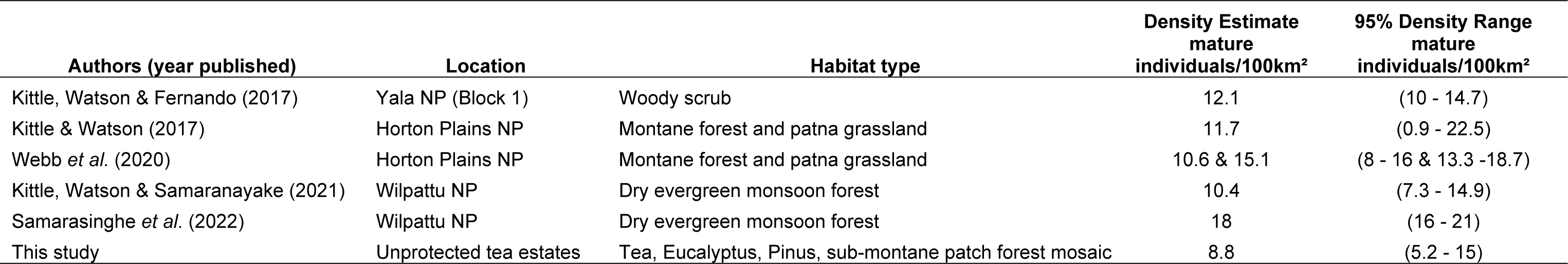
Leopard density (mature individuals/100 km²) comparison across published Sri Lankan studies that employed closed population, spatially explicit capture-recapture (R package *secr*) methods.

### Leopard site utilization

The model that best explained overall leopard site utilization was a co-existence model that included both human and dog RAI as well as their quadratic terms (Table 4). The coefficients for both quadratic terms were negative which is the only biologically plausible option [62] (Table S6) and indicates that leopard site use was maximized at intermediate levels of both human and dog abundance (*i.e.* both areas where human and dog occurrence were high and where human and dog occurrence were low, showed lower levels of leopard use). Model *R²* = 0.26.

**Table 4:**
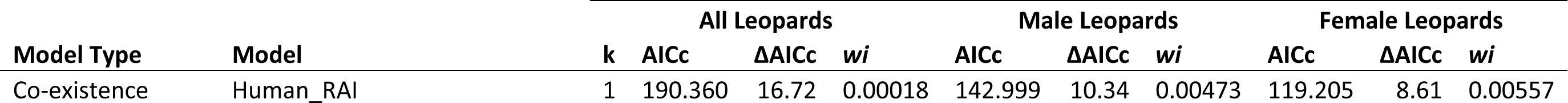
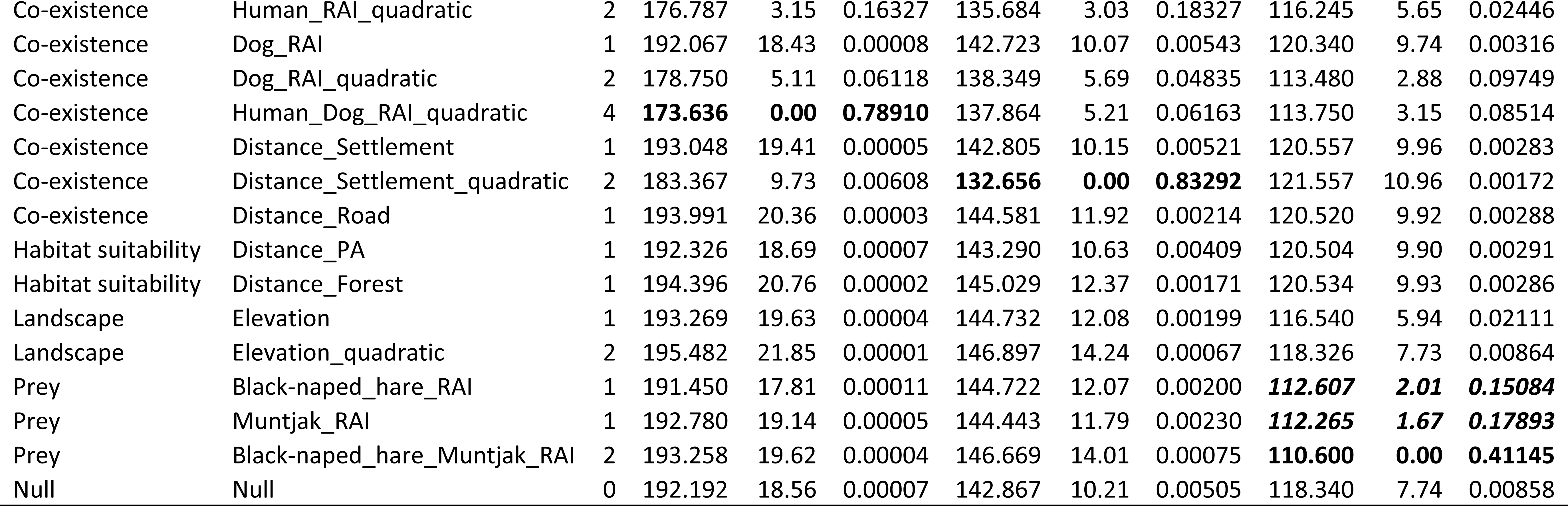
Comparison of all site utilization GLM models. Top models are in bold based on the lowest Akaike Information Criterion for small sample size values (AICc). If the difference between AICc values for the best model and a second model (ΔAICc) is < 2, those models are considered equivalent. Model weights (*w_i_*) are also shown.

Male leopard site utilization was again best explained by a co-existence model, this time the model incorporating both the distance to human settlement and its quadratic term (Table 4). Again, the quadratic term co-efficient was negative suggesting a preference for intermediate distance from human settlements (Table S5). Model *R²* = 0.19.

Female leopard site utilization was best explained by a prey model that incorporated both black-naped hare and muntjak RAI (Table 4). Both co-efficients were strongly positive indicating that female leopards were more likely to use sites with high occurrence of these prey species (Table S5). Model *R²* = 0.16.

### Leopard activity times

A total of 90 independent leopard observations were utilized for determining activity patterns from the unprotected tea estate landscape, with 183 observations from WNP [45] and 214 from GONP (Table S4). All three leopard populations were characterized as being active at dusk and nocturnally, with low diurnal activity (Table 5). Inter-site comparisons showed that there was no difference between the two protected areas, but leopards in the unprotected tea landscape were significantly less diurnal and more active around dusk than the protected area landscapes (Fig. 2; Table 6).

**Fig. 2:**
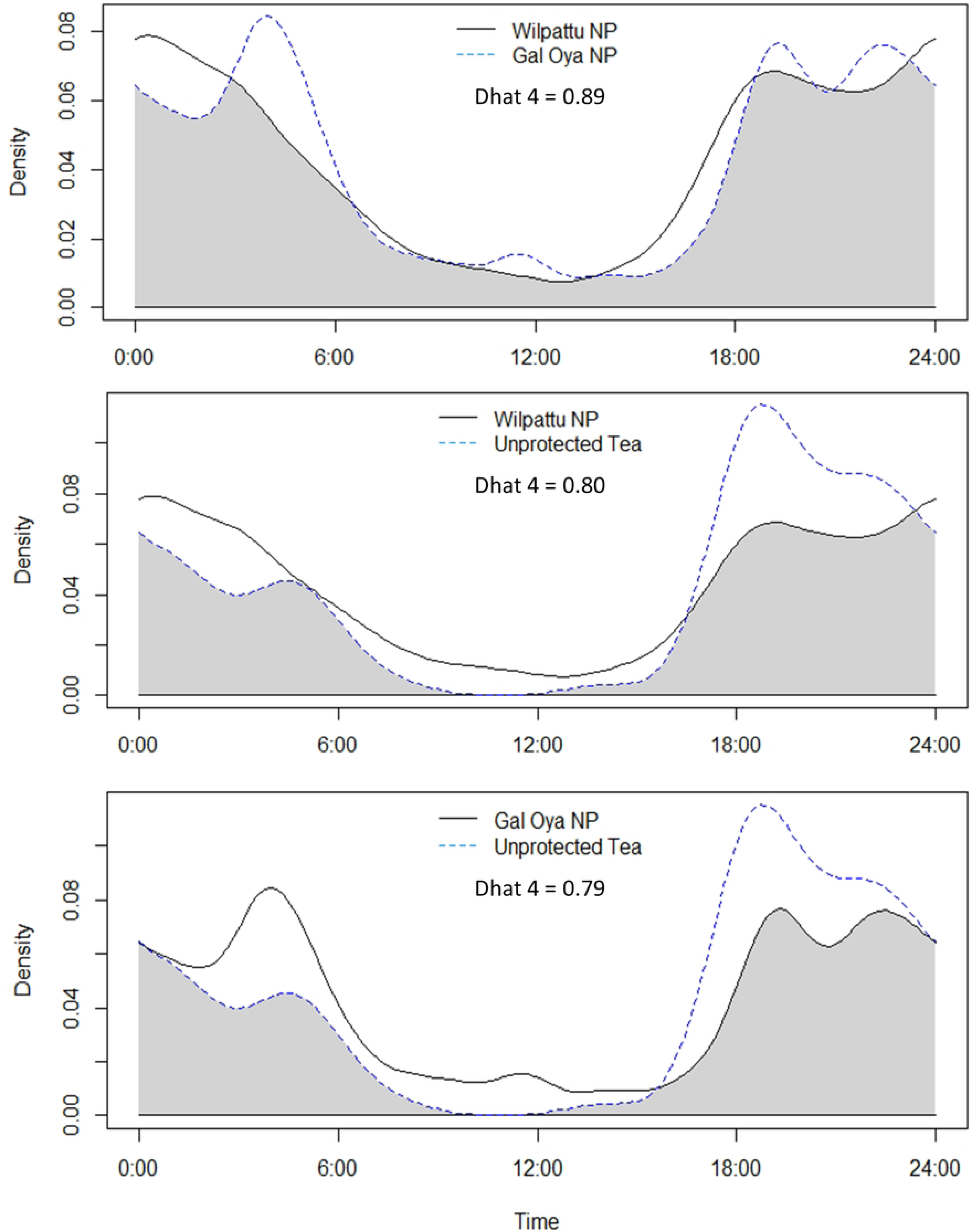
The fitted kernel density curves using default smoothing parameters (R package “overlap”) comparing two Sri Lankan Protected Areas (Wilpattu National Park and Gal Oya National Park; top), the unprotected tea estate landscape (this study) and Wilpattu NP (middle), and the unprotected tea estate landscape (this study) with Gal Oya NP (bottom). While leopards in all three study areas would be described as primarily active at dusk and nocturnally, the unprotected tea estate landscape leopards were significantly less active diurnally and more active at dusk than the two NP leopards.

**Table 5:** Activity period breakdown of leopards across three Sri Lankan study sites (2 National Parks and one unprotected landscape) based on remote camera photo-captures. Activity periods are Crepuscular either around dawn (Cr(Dawn)) or dusk (Cr(Dusk)), Diurnal and Nocturnal. The proportion of total observations during each period is shown (% Obs) as well as the Manly selection index (*w_i_*). When *w_i_* > 1 selection for the time period is indicated, whereas when *w_i_* < 1 avoidance is indicated, with greater selection or avoidance suggested as *w_i_*approaches 2 or 0 respectively.

| Location | Cr(Dawn)<br>5:15 - 7:00 |  | Diurnal<br>7:00 - 17:15 |  | Cr(Dusk)<br>17:15 - 19:00 |  | Nocturnal<br>19:00 - 5:15 |  | Category |
| --- | --- | --- | --- | --- | --- | --- | --- | --- | --- |
| | % Obs | $w_i$ | % Obs | $w_i$ | % Obs | $w_i$ | % Obs | $w_i$ | |
| Wilpattu NP | 6.0 | 0.82 | 14.2 | 0.33 | 11.5 | 1.57 | 68.3 | 1.60 | Cr(Du)/N |
| Gal Oya NP | 6.0 | 0.83 | 13.0 | 0.30 | 8.8 | 1.21 | 72.1 | 1.69 | Cr(Du)/N |
| Unprotected Tea | 4.4 | 0.61 | 4.4 | 0.10 | 23.3 | 3.20 | 67.8 | 1.59 | Cr(Du)/N |

**Table 6:**
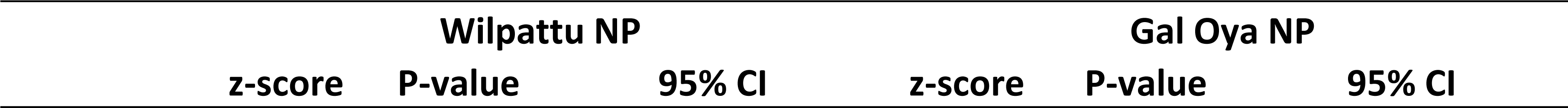
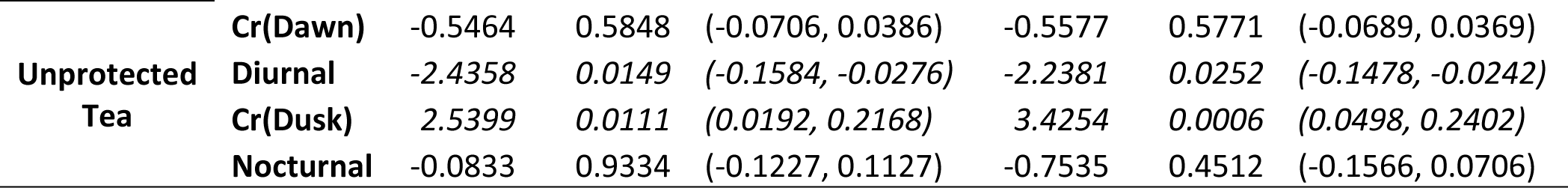
The comparison of independent proportions of leopard observations for each of the time periods (Diurnal, Nocturnal and Crepuscular dawn and dusk) between the unprotected tea landscape and the two National Parks (Wilpattu NP and Gal Oya NP). Two-sample, two-tailed Z-tests were employed with z-scores, P-values and 95% Confidence Intervals shown. When 95%CI do not overlap 0 it is considered strong evidence of difference between sites and is indicated here by italics.

## Discussion

### Leopard density

Leopard density in this unprotected, tea-estate dominated landscape was lower than what has been documented in three of Sri Lanka’s iconic, best protected National Parks – Yala, Wilpattu and Horton Plains - but this difference was far less stark than anticipated, with 95% CIs showing considerable overlap. This result, which suggests that protected areas might not automatically provide more suitable habitat for large carnivores, echoes results from Finland where the population densities of three of the four carnivore species in a nation-wide study - lynx (*Lynx lynx*), wolf (*Canis lupus*) and wolverine (*Gulo gulo*) - were unaffected by PA status [71]. These results suggest that the unprotected tea estate landscape, at least in this part of the Central Highlands, provides reasonable habitat suitability for the Sri Lankan leopard. This is likely due to several factors, including an available natural prey base [34,60], a rugged landscape that allows for refuge areas relatively inaccessible to humans [72], and the fact that the main crop being cultivated (tea) is a low, dense shrub that can provide good cover for movement [73]. Furthermore, although humans are widespread on the landscape and are active in the day, at nightfall people largely retreat back to their clustered communities leaving much of the region both dark and devoid of human presence. That leopards are renowned for their adaptability in human-dominated areas anyway [74], means that these key factors may be sufficient to allow a robust population here.

It is important to note that more recent surveys of protected area leopard populations in Sri Lanka have documented very high densities: 54.6/100 km² in Yala NP Block 1 [75] and 40/100 km² in Kumana NP which borders Yala NP to the NE [76]. However, these studies either did not use *secr* methodology [75] or did not comply with closed population assumptions in their density estimates [75,76]. Therefore, we felt unable to compare these estimates with those that used more standard methodology (Table 3). However, if these recent studies do reflect the on-ground situation and leopard densities of these NPs are increasing to such a degree, it represents a potentially worrying trend, as it suggests that these NPs might be becoming overcrowded, possibly due to increased habitat loss and associated anthropogenic pressures beyond PA boundaries [77]. Previous research in WNP has detected evidence of both potentially increased intra-specific competition amongst leopards [45] and potential inbreeding [54], both of which suggest populations being hemmed in by external forces.

While the adult sex ratio (ASR) of direct remote camera observations was female biased (37.5% M), this was even more marked (16.1% M) using the secr hybrid mixture model which estimates detection bias and sex-specific movement. Female biased ASRs have been detected regularly in large felids, and are to be expected given that most felids (including leopards) are polygynous with one male range overlapping the ranges of several females [78]. This pattern has been associated with the size of the study area, with larger study areas linked to increased female-biased ASR in both medium (∼50 kg) and large felids (> 100 kg) felids [79]. Although the current study area is not a bounded Protected Area, the remote camera survey conducted was representative of a much larger landscape - ∼2,400 km^2^ [34] - of broadly similar composition (*i.e.* a mosaic of tea cultivation, forest patches, plantation forests and communities), which aligns with the expectations of a female biased ASR.

Higher population density in large study areas has been similarly linked to increased female-biased ASR [79]. Male leopards have previously been shown to undergo density-dependent dispersal, with increased population density resulting in increased competition and higher rates of male emigration [80]. At the same time, when population density increases female leopards decrease their range sizes while male ranges remain stable, a factor which similarly results in a female-biased ASR [81]. The current survey described a surprisingly high adult leopard density in this unprotected, human-dominated landscape which, taken together with the landscape’s relatively broad extent, again aligns with expectations of female-biased ASR [79].

### Leopard site utilization

Leopards in the unprotected Central Highlands do not appear to be spatially avoiding humans, but to be highly cognizant of their presence, a factor which they seem to incorporate into space use decisions. Leopards were most likely to be found at intermediate levels of both human and domestic dog use, which perhaps suggests an avoidance of the most high-use areas [82] tempered by the ability to tolerate general human presence [11]. Dogs are unlikely to be a direct threat to leopards in this landscape – except potentially for small cubs - as most dogs are relatively small (∼ 15 kg) and don’t form packs here. Instead dogs are a prey source in this landscape, albeit comprising < 10 % of leopard diet and not being preferentially selected [60]. Leopards undoubtedly venture into tea estate communities and capture dogs on occasion [83], and despite being uncommon these incidents are unsurprisingly memorable to people who live in estate communities, which can result in an exaggerated perception of the frequency of this behavior. A similar situation is seen in the buffer zone of Sri Lanka’s Yala National Park, where the perception of leopard predation on livestock exceeds the actual toll that this predation takes [84]. In terms of promoting human-leopard co-existence, perception is more important than fact, and can lead to resentment, fear and retaliatory killing [85]. In India leopards are known to predominantly prey on domestic species in some regions where wild prey is limited or absent [11,86,87] which underscores the adaptability of leopards and suggests that maintaining a robust natural prey population in human-dominated landscapes is a key way to promote human-leopard co-existence.

Male leopards were seen to avoid human settlements which may be due to a perception of greater risk in proximity to humans [88]. Leopards within PAs have been observed to avoid PA boundaries in Sri Lanka [45] and Thailand [89] when those boundaries represent high levels of human activity. Female leopards were not seen to avoid human settlements, but instead were more likely to be found in areas of high natural prey – red muntjac and black-naped hare - availability. This is consistent with prey availability being a key factor underlying the size and location of female leopard home ranges [52,90]. Red muntjac are almost the ideal size for highly preferred leopard prey [61] so seeing leopards bias their space use towards areas frequented by this species is not surprising. Black-naped hare are smaller (∼2.5 kg) but are the most frequently predated species in the study area [60] and have very high levels of activity overlap with leopards here. Similar sized prey species, including the yellow-striped chevrotain (*Moschiola kathygre*) and even domestic dogs in this landscape, may be especially important prey species at key times for female leopards, as they regularly need to transport prey to den sites and/or dependent cubs [91,92].

### Leopard activity times

While leopards in this unprotected landscape showed similar general behavioral patterns as those within Sri Lankan PAs, in that they were predominantly active nocturnally and at dusk, tea estate leopards exhibited significantly less diurnal activity and were significantly more active around sunset than those within monitored PAs. It has been documented that wildlife in human-dominated landscapes often alter their behavior, typically expressed as increased nocturnal activity in order to avoid confrontations with people [28]. Here strict nocturnality did not increase in comparison to leopard populations in areas without humans, but daytime activity was reduced to the extent that it was almost negligible and compensation was made at nightfall, when a sharp spike in leopard activity was observed. This clear increase in activity around sunset further suggests a strategy of “lying low” during the day to avoid confrontations with people [89], with an increased need to move as soon as daytime ends. In unprotected areas outside Chitwan National Park in Nepal where, similar to this landscape, natural resource collection by humans is high during the day, temporal avoidance of humans was also especially pronounced [93].

This temporal partitioning is likely to be complemented by fine-scale spatial partitioning during periods of human activity. In Sri Lanka’s tea-estate dominated Central Highlands, the landscape is a mosaic of land use types with vast expanses of tea cultivation interspersed with pockets of natural forest, plantation forest and scrub/shrub areas where previous tea cultivation has been discontinued. It is in these areas that leopards appear to spend the daytime hours before emerging to roam the extensive network of tea estate roads and walking paths – used by estate workers and community members during the day - that traverse the landscape. Cultivated tea fields can provide adequate cover for leopards to move through, but given the regular daytime presence of estate workers in these areas and the relatively low frequency with which leopards are sighted here, it is unlikely that they are extensively used as daytime resting places. However, small cubs have been found on several occasions within the tea bushes, left there by temporarily absent mothers, which suggests that leopards do sometimes utilize the tea fields in this way.

Although temporal partitioning can be an effective way to ensure co-existence in shared landscapes (e.g. [94]) particularly for persecuted species or predators [95], wire snares, which remain on the landscape even when people have left, transcend this adaptation by de-coupling the threat from the presence of humans. This threat is particularly prevalent in Sri Lanka’s unprotected Central Highlands and is the leading known cause of leopard mortality in this region [35]. Widespread snaring can have a profound impact on mammalian biodiversity [96,97], including apex predators whether they are targeted [98] or not [99]. While snares in the Central Highlands of Sri Lanka typically are not set to target leopards, being unselective they nevertheless take a toll [35]. In order to ensure the long-term viability of leopards in this unprotected landscape it is essential to curb the use of wire snares.

## Conclusion

This unprotected, hill country leopard population exhibits a density lower than Sri Lanka’s PAs for which comparable data are available, however the difference was considerably smaller than anticipated. Furthermore, the 3 compared PAs – Yala (Block 1), Wilpattu and Horton Plains - are Sri Lanka’s most iconic National Parks, all of which are well known as leopard “hotspots”. This suggests that the tea landscape of the southern Central Highlands is conducive for leopard presence despite the widespread and high level of human activity found here. Spatial partitioning is not how this relatively robust leopard population negotiates potential anthropogenic impacts, although results suggest that leopards here do consider human presence when making space use decisions. Instead, it appears that temporal partitioning, with decreased leopard activity in the day compensated by increased activity around dusk, is the behavioural adaptation which underlies human-leopard co-existence here.

To summarize the findings of what is the first comprehensive study of leopard density and behaviour outside of the island’s protected areas, leopards appear to have adapted well to this unprotected, human-dominated landscape with temporal partitioning a clear behavioral adaptation undertaken by them to reduce direct human threats and essentially foster human-leopard co-existence. From a management perspective, to ensure that the island’s leopard population remains robust in the long term, the leopards in this unique, vulnerable region of Sri Lanka require ongoing monitoring and to be subject to conservation initiatives aimed at ensuring their long term viability. These include the maintenance of existing habitat patches and corridors that allow, at least, for the present level of connectivity. Key to the implementation of conservation initiatives is the need to maintain or strengthen present levels of human-leopard co-existence. To do this it is imperative to 1) maintain current temporal patterns (limited nighttime use by humans) to avoid off-setting leopard behavior adaptions, 2) ensure that the region’s sizeable natural prey base remains stable as this keeps leopards from switching to a diet more reliant on domestic species, and 3) minimize – or ideally, eliminate - the use of snares in this region, as these can transcend behavioural adaptations and render this leopard population highly vulnerable.

## Acknowledgements

Sincere appreciation is given to all of the Regional Plantation Companies (RPCs) and private landholders that allowed us to conduct this study on their estate lands. This includes Maskeliya Plantations PLC, Bogawantalawa Tea Estates PLC, Madulsima Plantations PLC, Dilmah/MJF Tea Gardens, Horana Plantations PLC, as well as Mr. Balendran at Kelani-Braema private tea estate. Resplendent Ceylon has been supporting this work since its inception and generously provided a bungalow from which to conduct this research. Our field team of Emad Sangani and Riahn Pieris, ably assisted by Maya Sithunayake and Nimalka Sanjeewani, were essential to keeping the cameras operational. The Sri Lanka Department of Wildlife Conservation (DWC) gave permission to conduct this study. Thanks also to CERZA Conservation, Z-Gap and OLU Tropical Water for support.

## Supporting Information captions

**Table S1:** Remote camera location coordinates and running durations for this study.

**Table S2:** Remote camera array buffer estimation. The “check.buffer” function of the R “secr” package was used to determine the buffer size, which was selected when the Log Likelihood values stabilized at 3 decimal places.

**Table S3:** Final spatial utilization model variables.

**Table S4:** Remote camera location coordinates and running durations for Gal Oya NP.

**Table S5:** Monthly sunrise and sunset times for the study area.

**Table S6:** Coefficient values, standard errors, z-values and P-values for variables in all top spatial models. P-values < 0.1 are italicized, with p-values < 0.05 italicized and in bold, and p-values < 0.01 italicized, bold and with an asterisk.

